# Alternative 3’ Polyadenylation Responses to Acute Ethanol Exposure Differ Between *Drosophila* Populations

**DOI:** 10.64898/2026.03.05.709877

**Authors:** George Boateng-Sarfo, Seungjae Lee, Eric C. Lai, Sarah Signor

## Abstract

Alternative polyadenylation is a pervasive post-transcriptional mechanism that generates mRNA isoforms with distinct 3’ untranslated regions (3’ UTRs), which can influence transcript stability, localization and translation. Ethanol is an environmental stressor for *Drosophila melanogaster*, and natural populations show distinct responses to ethanol exposure, suggesting that the genetic background can modulate phenotypic response. Yet, how the natural genetic variation modulates ethanol-responsive APA remains understudied. We profiled 3’ end isoforms following acute ethanol exposure in three French (cosmopolitan) and three Zambian (ancestral range) *D. melanogaster* genotypes and identified numerous significant APA events across all genotypes. Ethanol-induced APA was widespread but highly genotype - and population - dependent: the French genotypes showed an enrichment of proximal shifts (3’ UTR shortening), while Zambian genotypes were biased toward distal shifts (3’ UTR lengthening). Most of the ethanol-responsive APA events were private to a single population, with relatively few genes exhibiting conserved directional changes across both backgrounds. We also classify genes showing opposite-direction shifts between populations, and others with opposite-direction shifts between various classes of APA. For two-site events, inter-poly(A) site distance did not predict response direction; however, spacing strongly predicted the magnitude of remodeling, with larger separations enabling greater absolute 3’ UTR length changes. Our findings support a model in which ethanol-responsive 3’ end regulation is a flexible genetically contingent, and that population-specific APA remodeling represent a plausible molecular substrate contributing to adaptation.

## Introduction

Alternative mRNA processing, such as alternative splicing and alternative polyadenylation, have been demonstrated to be important for macro-evolutionary trends such as organismal complexity. For example, the number of coding genes and the length of their coding sequence has remained relatively stable between worms and humans^1–3^. However, the number of genes that produce alternative 3’ UTRs has doubled, and the length of those UTRs has increased 8.5x^4,5^. Alternative 3’ UTRs mediate *cis*-regulatory interactions with other proteins which affects stability, localization, and translational efficiency. This is important for cell-type specific gene expression and organismal development.^4–12^

However, the role of alternative splicing and polyadenylation of mRNA in micro-evolutionary processes is unclear. One area where alternative polyadenylation may be relevant is during stress responses and adaptation to changing environmental conditions. For example, in some stress genes in plants non-functional transcripts are produced in the absence of stress. Also, during stress response they are rapidly converted to functional transcripts, avoiding the negative effect on growth when they are not needed but also avoiding the time necessary for pre-mRNA accumulation and transcription. While the role of gene expression overall in stress and adaptation is well established, we currently know very little about the role of alternative mRNA processing.

Gene expression differences in response to ethanol have been intensely studied in *D. melanogaster*^13–29^. However, alternative mRNA processing including alternative polyadenylation has been paid little attention^16,30^. Ethanol in the decaying fruit that *D. melanogaster* regularly feeds and oviposites in can reach concentrations of 6-7%^31,32^. Ethanol is an environmental stressor that is avoided by most other *Drosophila* species^33^. Its sister species, *D. simulans*, has about 4x lower the tolerant to ethanol and avoids ethanol rich substrates^13,32,34^. *D. melanogaster* became adapted to ethanol as it spread out of Africa and became a human commensal. The ancestral range of *D. melanogaster* is thought to be around Zambia in sub-Saharan Africa^42^. Demographic models suggest that *D. melanogaster* began its expansion out of Africa 10,000 years ago and colonized Europe approximately 2,000 years ago^35,36^. These recently separated populations differ in their ethanol resistance with the Zambian populations having low resistance to ethanol and the populations from colder latitudes such as France having higher ethanol resistance^37^. Thus, this represents a recently evolved difference between these populations.

Given that ethanol is a common environmental stressor that *Drosophila* encounters frequently and alternative polyadenylation is a plausible route to adaptation, we profiled 3’ UTR poly(A) site usage in French and Zambian population. We found that ethanol induces widespread but heterogenous APA changes in both populations. Additionally, we showed that these patterns point to APA as a flexible, genotype-dependent mechanism by which ethanol can modulate mRNA fate beyond gene expression. Here, we test whether ethanol-responsive APA differs systematically between populations that vary in ethanol tolerance, and whether the direction and magnitude of APA shifts are driven primarily by population background, genotype, or poly(A) site architecture.

## Materials and Methods

### Fly strains

Fly strains were generously provided by John Pool (UW-Madison) and originated either from France or Zambia^38,39^. The samples from Zambia were previously confirmed to be the most diverse among all the globally sampled strains, with minimal non-African admixture, suggesting they come from the ancestral range of *D. melanogaster*^38,39^. We used three strains from Zambia and France respectively for three replicates of each condition (3 replicates x 6 genotypes x 2 conditions).

### Fly collection

To minimize variation due to non-focal effects, all collection populations were set up with 10 one day old individuals of each sex from a particular genotype. Flies were reared on a standard Bloomington medium at 25°C with a 12-h light/12-h dark cycle. After 8-9 days the vials were cleared and F1 males were collected for ethanol exposure assays and RNA-seq. 30 mated males were collected within three days of one another and subjected to assays within 3-5 days.

### Experimental Setup

The flies were sedated by exposing them to cold for 10 mins and placed in the behavioral chambers with a paintbrush. Each chamber contained 5 ml of standard grapefruit fly media or media in which 15% of the water had been replaced by ethanol. They were allowed to acclimate for 10 min prior to timing the 30-minute exposure. This acclimation period is standard for behavioral assays, as it is long enough for the initial startle response to ethanol to have concluded, however here it was included to standardize data with past observations^16,30,40–43^. The assays were conducted within a 2-hour window after dawn, the period in which the flies are most active^44–46^. Replicates were conducted randomly across days under standardized conditions (25°C, 70% humidity). At the conclusion of the assay the flies were flash frozen in liquid nitrogen. Frozen nested sieves were used to separate their bodies from their heads, limbs, and wings. Bodies were collected for sequencing.

### Behavioral assays

Differences in the response to ethanol has been previously observed in African and cosmopolitan *D. melanogaster*^47–50^. However, we wanted to confirm that African populations had a lower tolerance for ethanol. A lower tolerance for ethanol would be indicated by a quicker progression through the euphoric stage of alcohol exposure to sedation^16,51^. Approximately 30 male flies from a single genotype were sedated in a refrigerator for 10 minutes and then placed in a petri dish with 5 ml of grapefruit media in which 15% of the water had been replaced with ethanol. The petri dish was bisected with a black line, and every minute the flies were observed for 20 seconds. The number of flies that crossed the black line were recorded as a proxy for activity level, indicating that the flies were not sedated if they crossed the black line. This was done for ten minutes. Three replicates for each of the three genotypes was performed (Supplementary file 1).

### 3’ UTR sequencing

RNA was extracted from the bodies of 30 male flies using the NucleoZol one phase RNA purification kit (Macherey-Nagel). Library preparation and sequencing was performed by Lexogen (Vienna, Austria) using the QuantSeq 3’ mRNA-Seq Library Prep Kit REV. This kit generates illumina compatible libraries of sequences at the 3’ end of polyadenylated RNA. The libraries were barcoded and pooled, and 2 million reads were generated per library on the illumina NextSeq. The data was demultiplexed prior to delivery.

### 3’ UTR data analysis

The 3’seq reads were subject to quality filtering and adapter trimming using fastp^52^. To quantify alternative polyadenylation (APA) dynamics, we used the software package LABRAT^53^, a transcriptome level APA analysis tool that infers APA site usage across experimental conditions. We specified the 3pseq libtype which is appropriate for 3’ sequencing rather than RNA-seq. Briefly, transcripts were mapped to the *D. melanogaster* transcriptome (dmel-all-transcript-r6.64.fasta)^54^ and quantified using *salmon* ^55^. The output files from salmon were used by LABRAT to calculate Ψ values for each gene across all genotypes. Thus, the Ψ values represent the proportion of a gene’s isoforms that utilizes the distal APA relative to the total transcript output of that gene. As such, the Ψ values range from 0 to 1, where values closer to 0 indicate proximal/upstream usage of APA sites while values closer to 1 indicate downstream/distal APA site usage (Supplementary file 2 and Supplementary file 3).

LABRAT assigns genes to three APA types based on the structural context of their APA sites Figure 2 A-C. These are tandem untranslated region (tandem UTR) for APA events that occur within the same 3’UTR (Figure 2A); alternative untranslated region (alternative UTR) for APA events that are located in different terminal exons (Figure 2B); and mixed untranslated region (mixed UTR), where the transcripts models suggest overlapping usage of both types (Figure 2C).

Differential APA usage for each category of UTR usage mentioned above was assessed by computing Ψ values as defined by LABRAT: Ψ(ER) - Ψ(NR) where ER refers to the ethanol treated samples and NR to the untreated samples. The results reflect the relative change in distal sites usage between the conditions. Hence, positive Ψ signifies increased distal APA usage in the ethanol-treated samples while a negative Ψ denotes an increase in proximal site usage in response to ethanol exposure. For downstream analyses, only genes with statistically significant APA changes (*p* ≤0.05) and possessing at least two APA sites separated by ≥ 25 nucleotides were retained (25 nt is defined as the minimum separation by LABRAT)^53^. *p*-values were corrected for multiple testing using the False Discovery Rate^60^. This retains only high confidence APA events with meaningful isoform diversity. Unless otherwise stated, all significant APA events refer to FDR-adjusted p-value < 0.05.

We then tested whether the direction of APA shifts (distal vs proximal) differed by population, genotype, and APA class. Rather than fitting gene-level models, we summarized the significant genes into group-level counts of distal and proximal shifting events for each genotype within and each population (stratified by APA class). We then modeled the proportion of distal shifts with a binomial generalized model (logit link), of the form cbind (distal, proximal) ∼ APA type × population + population:genotype. This allowed us to group genotype within population, so they are compared within their populations and predictors do not overlap. Then we evaluated the significance of the terms using a likelihood-ratio tests, comparing the deviance of the full model to that of reduced models and using the chi-square (χ²) with appropriate distribution and degrees of freedom (Supplementary file 4).

We then obtained model-adjusted distal estimates and 95% confidence intervals for each genotype within each population using marginal means from the fitted generalized linear model. These estimates were averaged over genotype (alternative UTR, tandem UTR or mixed UTR) with proportional weights, so categories with more contributing genes have greater influence. Where contrasts were of interest, we expressed differences as odds ratios relative to a baseline genotype within each population and reported confidence intervals and multiple-testing-adjusted *p-values* for those comparisons (Supplementary file 4).

## Results

### Behavioral assays

**A)** *D. melanogaster* from Zambia had a significantly lower tolerance to ethanol than French *D. melanogaster* (Figure 1D, two tailed t-test *p* < .0001)^57^. French flies take on average twice as long to become intoxicated as Zambian flies (Figure 1E). As was previously reported, this confirms the expected difference in ethanol tolerance between African and cosmopolitan *D. melanogaster*^57^. Consistent with prior work, French genotypes show higher ethanol tolerance than Zambian genotypes, providing a phenotypic context for interpreting population differences in ethanol-responsive APA. These results confirm that the same genotypes used for 3’UTR profiling differ in ethanol sensitivity at the behavioral level, establishing a phenotypic context for interpreting molecular difference in APA regulation.

**Figure 1:**
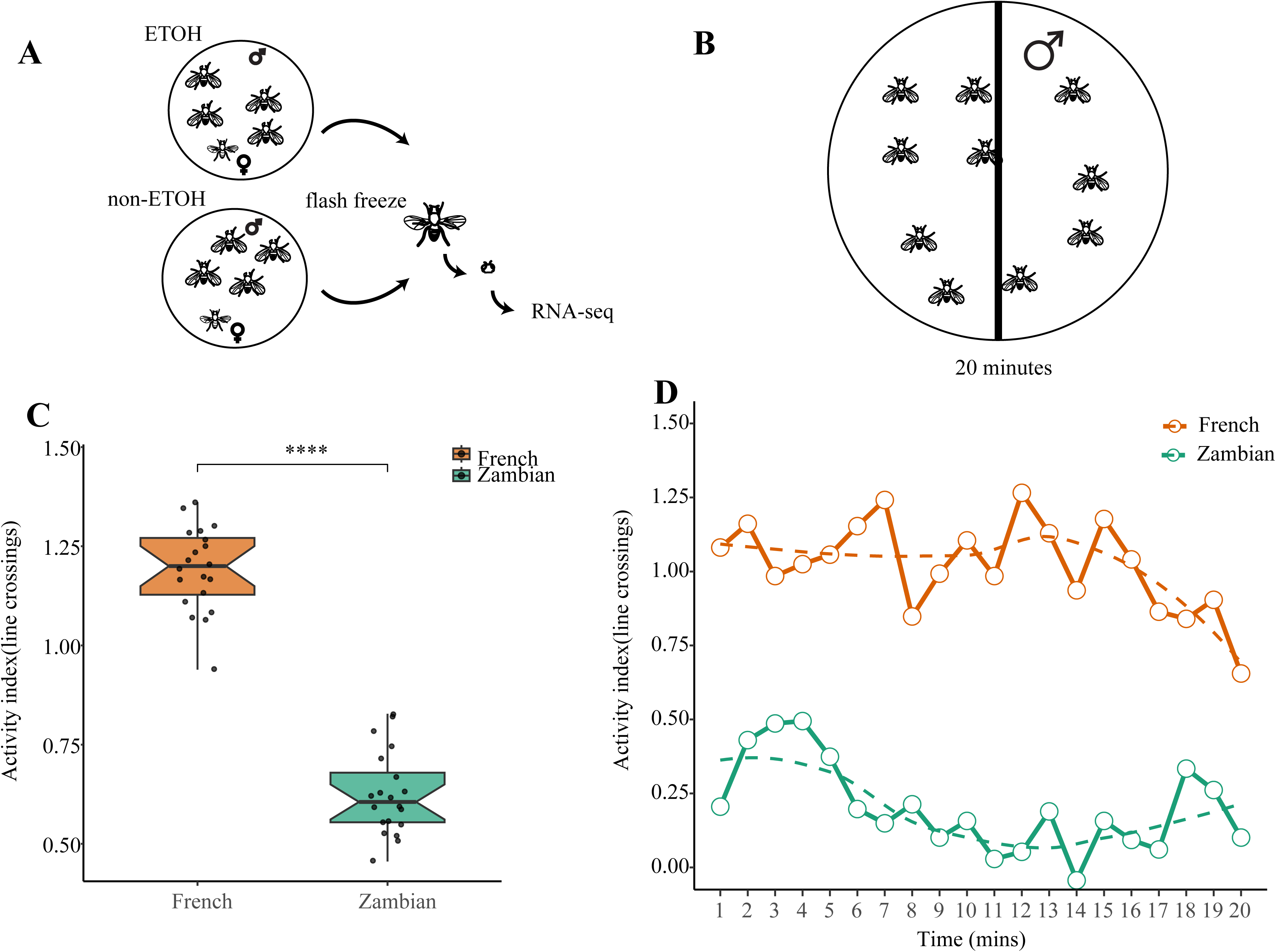
Experimental design and ethanol sedation across the French and Zambian population. **A)** An illustration of the environment that each *Drosophila* male was exposed to during the experiment. **B)** An example of the behavioral setup used to confirm that the two populations of *D. melanogaster* have different responses to ethanol. Every minute for twenty minutes, the number of flies that crossed the midline of the plate was recorded. The substrate contains 15% ethanol. **C)** The proportion of flies sedated after ethanol exposure showing that the French population had a higher tolerance relative to the Zambian. ****= p-value ≤ 0.0001. **D)** The times analysis illustrating that the French population maintained a higher and more stable sedation level over time, while the Zambian population have lower proportion and more variable.

**Figure 2:**
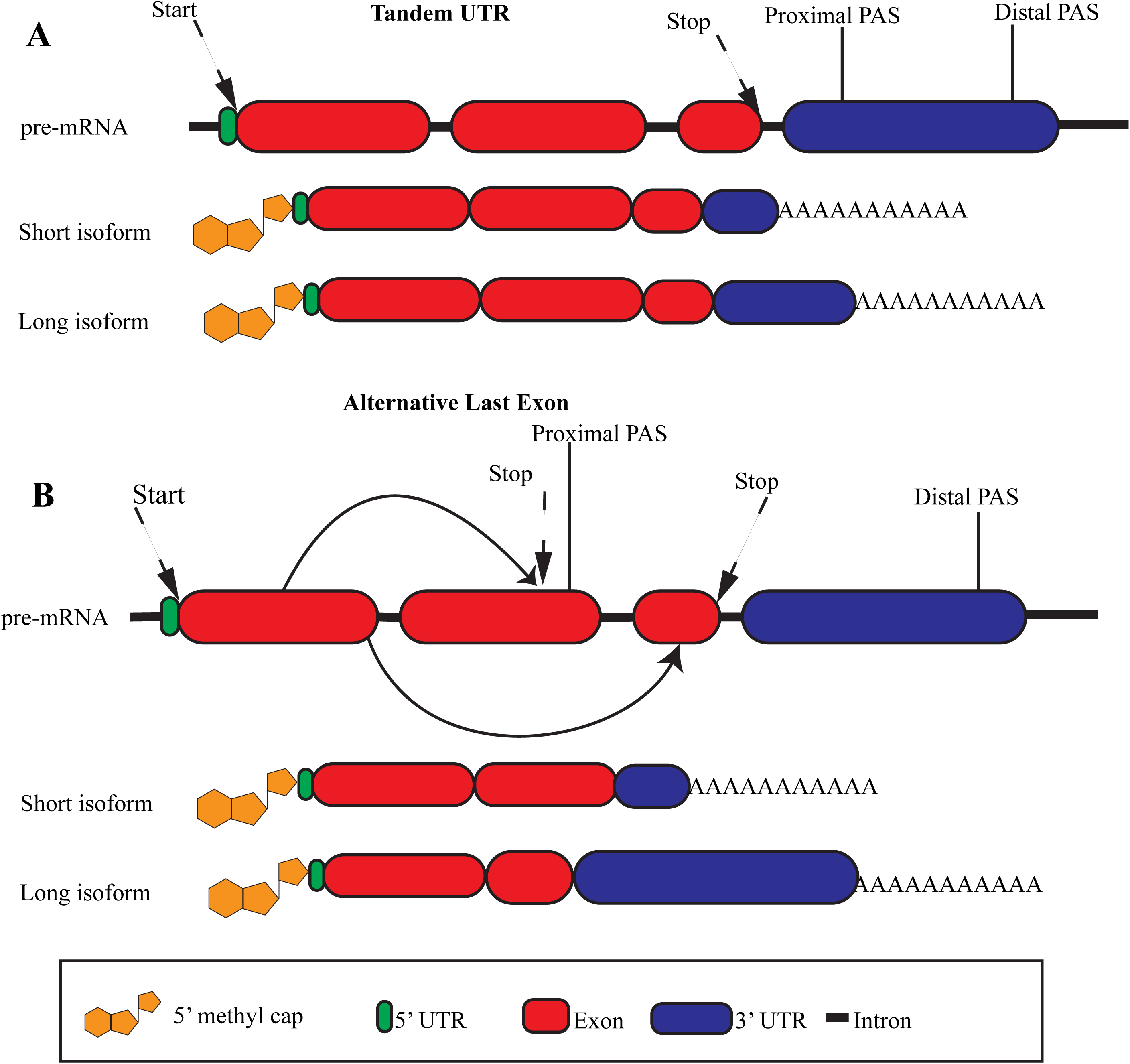
Alternative polyadenylation mechanisms of generating 3’ transcript isoforms. **A)** Schematic of tandem 3’ UTR APA illustrating the transcription start and end sites, and proximal and distal PAS. In tandem APA, the 3’ terminus can either be cleaved upstream (Proximal) or downstream (Distal), thus, resulting in different 3’ ends. **B)** Schematic of alternative last exon, in which differential splicing selects distinct terminal exons that terminate at a cleavage site, leading to different 3’ ends. Also showing is the Proximal PAS that can lead to different termination coding sequence.

### 3’ UTR changes in response to ethanol

To determine whether ethanol induces population-specific 3’ UTR remodeling across genotype, we quantified APA shifts in the French and Zambian *D. melanogaster*. In total of 319 APA events (*p* ≤ 0.05) were detected across all genotypes from the two populations. Of these, 193 were found in French genotypes (*FR109*, *FR112* and *FR113*), and 126 APA events were found in the Zambian genotypes (*ZI274*, *ZI31*, *ZI418*). Notably, some of these APA events were shared between the genotypes from both populations. The distribution of APA events was also categorized into alternative UTR, tandem UTR, and mixed UTR based on the polyadenylation sites (Figure 3A).

**Figure 3:**
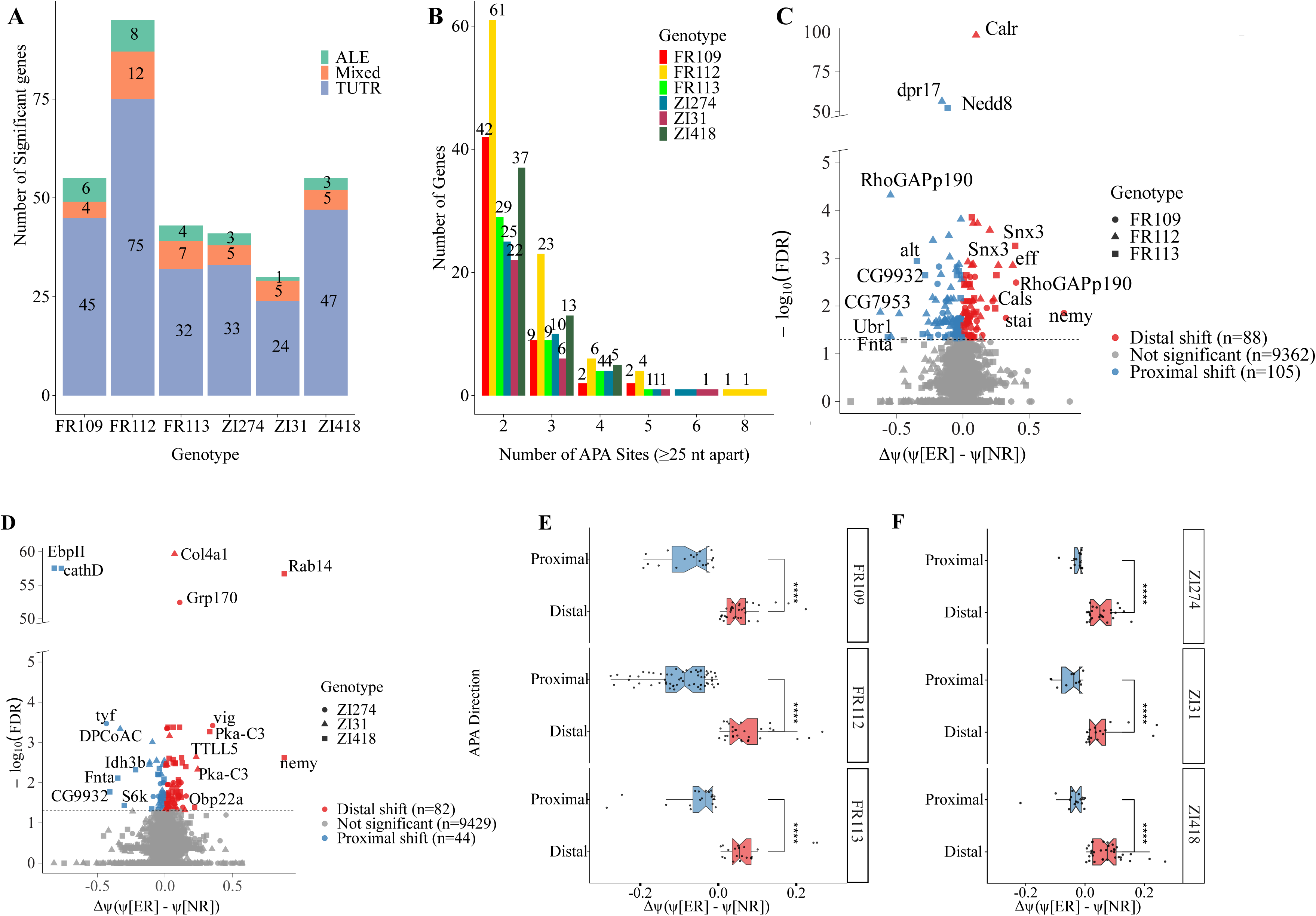
APA distribution in French and Zambian populations after ethanol exposure. **A)** The bar plot displays the number and distribution of APA events across distinct genotypes and populations. Each bar corresponds to the APA classification profile of a single genotype. Tandem Untranslated Region (TUTR) events are shown in purple, Mixed APA events in brown, and Alternative Last Exon (ALE) events in green. **B)** A bar plot summarizing the number of APA sites detected per gene across genotypes. Each genotype is color-coded to facilitate comparison. The red, gold, and green bars correspond to the *FR109, FR112,* and *FR113* samples, while cyan, magenta, and olive represent *ZI274*, *ZI31*, and *ZI418*, respectively. **C**)A scatter plot showing individual genes with either distal or proximal 3’UTRs in the French population. **D**) A similar scatter plot illustrates APA shifts in the Zambian population. The x-axis denotes the ΔΨ values (ethanol-treated - control), while the y-axis represents the -log10(FDR), indicating statistical significance. The grey dashed line marks FDR cutoff at 0.05, denoting the threshold for identifying significant regulated APA events. In both panels, blue colored shapes represent Proximal APA shifts while the red colored shapes represent Distal shift. In addition, the circular shape represents *FR109/ZI274*, triangle and square shapes represent *FR112/ZI31* and *FR113/ZI418* respectively. **E)** A boxplot showing the central tendency, variability, and range of ΔΨ values for significant distal and proximal APA events in the French population, as identified by LABRAT. **F)** A corresponding notched boxplot for the Zambian population is shown. In both plots, blue indicates proximal APA usage, while red represents distal APA usage. The blue colored box plots represent proximal APA shifts while the red colored shapes represent Distal shift.

Within the French population, the *FR109* genotype contributed 55 of events, including 45 tandem UTR events, 4 mixed UTR events, and 6 alternative UTR events. The most dynamic genotype was *FR112*, with 95 APA events, including 75 tandem UTR events, 12 mixed UTR events, and 8 alternative UTR events. In addition, we observed a more modest response in the *FR113* genotype, with 43 calls comprising 32 tandem UTR events, 7 mixed UTR events, and 4 alternative UTR events. Despite differences in the total number of events, all three French genotypes maintained a similar proportion of differences in tandem UTRs suggesting tandem UTR switching as their dominant APA mechanism.

Among the Zambian population, *ZI274* accounted for 41 events 33 tandem UTR events, 5 mixed UTR events, and 3 alternative UTR events. *ZI31* exhibited 30 events composed of 24 tandem UTR events, 5 mixed UTR events, and 1 alternative UTR event. *ZI418* stood out as the most plastic genotype, with 55 events which includes 47 tandem UTR events, 5 mixed UTR events, and 3 alternative UTR events. Similarly, the Zambian genotypes all showed tandem UTR changes as the predominate mode of 3’ UTR regulation. Within each population, most significant events were again, the tandem UTR type across genotypes. The number of ethanol-responsive APA events varied by genotype, indicating substantial genotype-dependent variation in 3’ UTR remodeling.

### Number of poly(A) sites in actively regulated 3’ UTR genes

Given the broad ethanol-responsive APA signal, we next asked how much poly(A) sites underlies these regulated events - especially if responsive genes typically harbor only two sites or contain multiple APA sites that could support more diverse 3’ UTR outcomes (Figure 3B). We noticed that the majority of APA regulated genes harbor two or more poly(A) sites. In fact, 132 genes in the French population which includes 42 genes in *FR109*, 61 in *FR112*, and 29 in *FR113* had exactly two APA sites. Similarly, *FR112* exhibited a richer multi-site distribution than the other French genotypes (23 genes with 3 sites, 6 with 4, 4 with 5, and one extreme case with 8), indicating greater APA complexity in this genotype. Examples of genes with two APA sites include *stathmin*, *Dorsal-related immunity factor*, and *nervana 3*. Whereas genes with 3 APA sites include *Imaginal disc growth factor 3*, *poly(A)-binding protein*, and *Na+,K+-ATPase alpha subunit*. Four-site cases include *Calmodulin*, *held out wings*, and *C-terminal Binding Protein*. We also observed 5-site genes such as *orb2*, *Tousled-like kinase* and *oo18 RNA-binding protein 2*, and an extreme 8-site case in *hu li tai shao*. Interestingly, *orb2* is known to bind to the 3’ UTR of mRNA and different isoforms have contrasting effects on translation^58^. We noticed that *Calmodulin* stood out as significant gene among all three French genotypes.

Among Zambian genotypes, 25 genes were identified in *ZI274*, 22 in *ZI31*, and *37* in *ZI418* with exactly two APA sites. These *include NADH dehydrogenase [ubiquinone] 1 subunit C2 (Complex I subunit B14.5b), mitochondrial ribosomal protein L28 (39S ribosomal protein L28, mitochondrial), Daisho1 (antimicrobial/immune effector peptide), ATP-binding cassette subfamily D member 1 (peroxisomal ABC transporter), no extended memory (cytochrome b561 family protein), and eukaryotic translation initiation factor 3 subunit A*. In addition, a few genes demonstrated greater APA complexity, harboring three or more alternative poly(A) sites. *ZI418* has the most multi APA site among the Zambian genotypes (13 genes with 3 sites, 5 with 4, 4 with 5), *ZI274* is intermediate (10 with 3, 4 with 4, 1 with 5, and 1 with 6), and *ZI31* has the fewest multi-site cases (6 with 3, 1 with 4, 1 with 5, and 1 with 6).

Additionally, we noticed that APA class differed by population (Figure 3C-D). For instance, the French genotypes showed more proximal increasing events (Figure 3C), whereas the Zambian showed more distal increasing (Figure 3D). A binomial generalized linear model (logit; outcome = distal vs proximal) revealed that *FR109* used distal usage 65.2% (95% CI 51.7-76.6), *FR112* was 32.6% (24.0-42.7) indicating a proximal shift, and *FR113* was intermediate at 49% (34.5-63.7). In the Zambian population, the genotypes were uniform biased towards distal APA usage (*ZI274* 65.9% (50.2–78.8), *ZI31* 59.2% (41.0–75.3), *ZI418* 66.7% (53.1–78.0)) implying nuance differences between the genotypes.

We then asked whether ethanol alters effect size at the event level (Figure 3E-F). Among the French genotypes, we noticed that proximal increasing events were broader in the *FR109* and *FR112*, while *FR113* showed a subtle shift (Figure 3E). In the Zambian genotypes, the dispersion flips. Thus, distal increasing events are broader, particularly in *Z1418* and *Z131* while modest in *Z1274*. Taken together the data suggests that ethanol-responsive APA predominantly acts on genes that already have multiple annotated poly(A) sites, and the direction of ethanol-induced shifts differ by population. Thus, the Zambian genotypes show stronger distal-increasing APA shifts, while French genotypes show modest, genotypes specific shift with no consistent distal bias.

### The distance between poly(A) sites and its effect on 3’ UTR length

We then sought to determine if the distance between APA sites helps explain either the direction or magnitude of the ethanol-induced 3’ UTR remodeling. We focused our analysis our analysis on two-site events and summarized inter-poly(A) distances irrespective of direction of shift (Figure 4A). Across all six genotypes, the alternative sites were typically separated by hundreds of nucleotides and on some occasions kilobase scale gaps. The French genotypes span broader ranges, especially in *FR109/FR112*, whereas *FR113* is comparatively narrow. On the other hand, in the Zambian population, the distance is generally more concentrated (most notably *ZI418*), with *ZI274*/*ZI31* showing only occasional long-range outliers. These patterns indicate that genotype/population background modulates typical 3’ end spacing.

**Figure 4:**
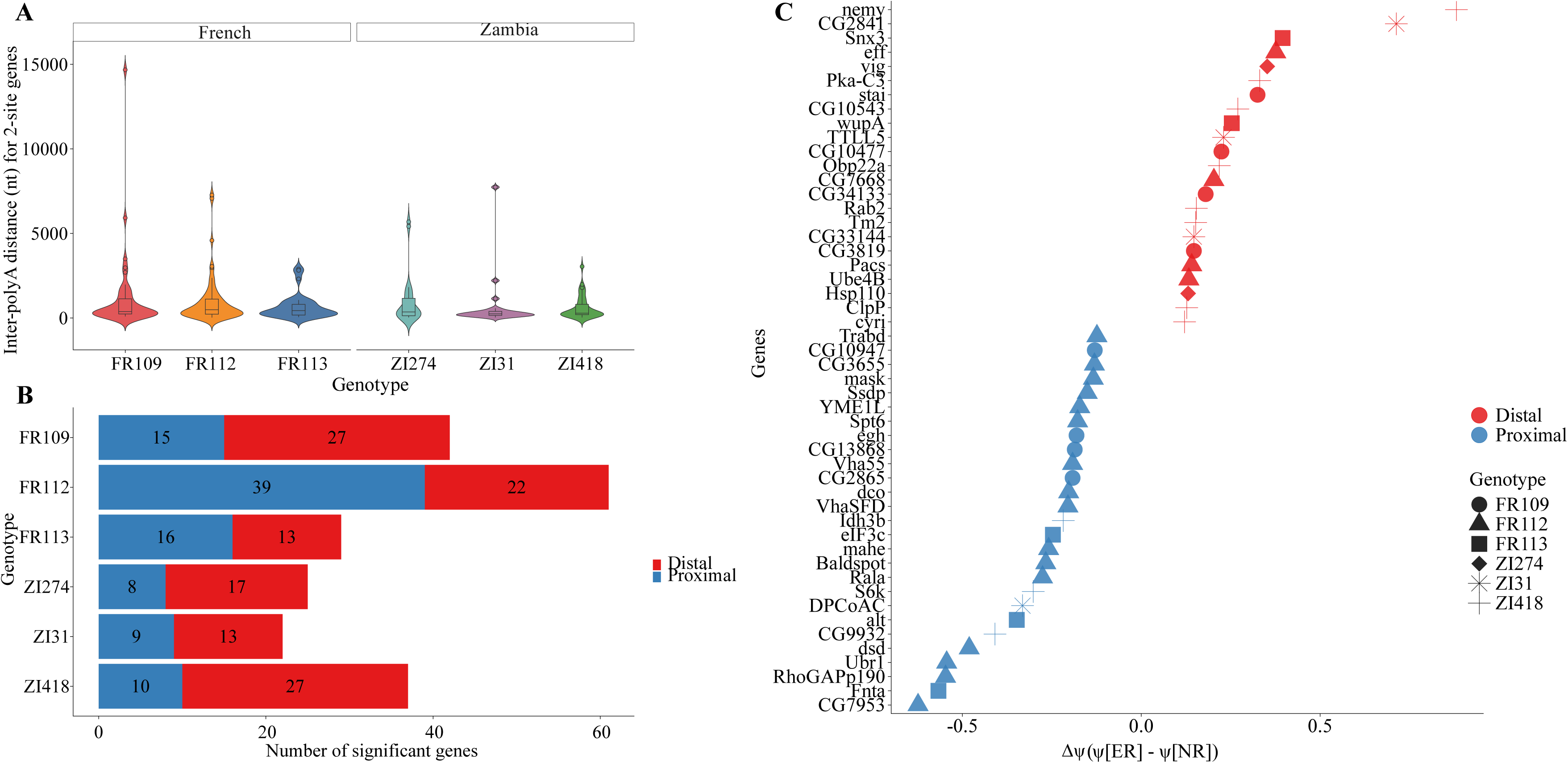
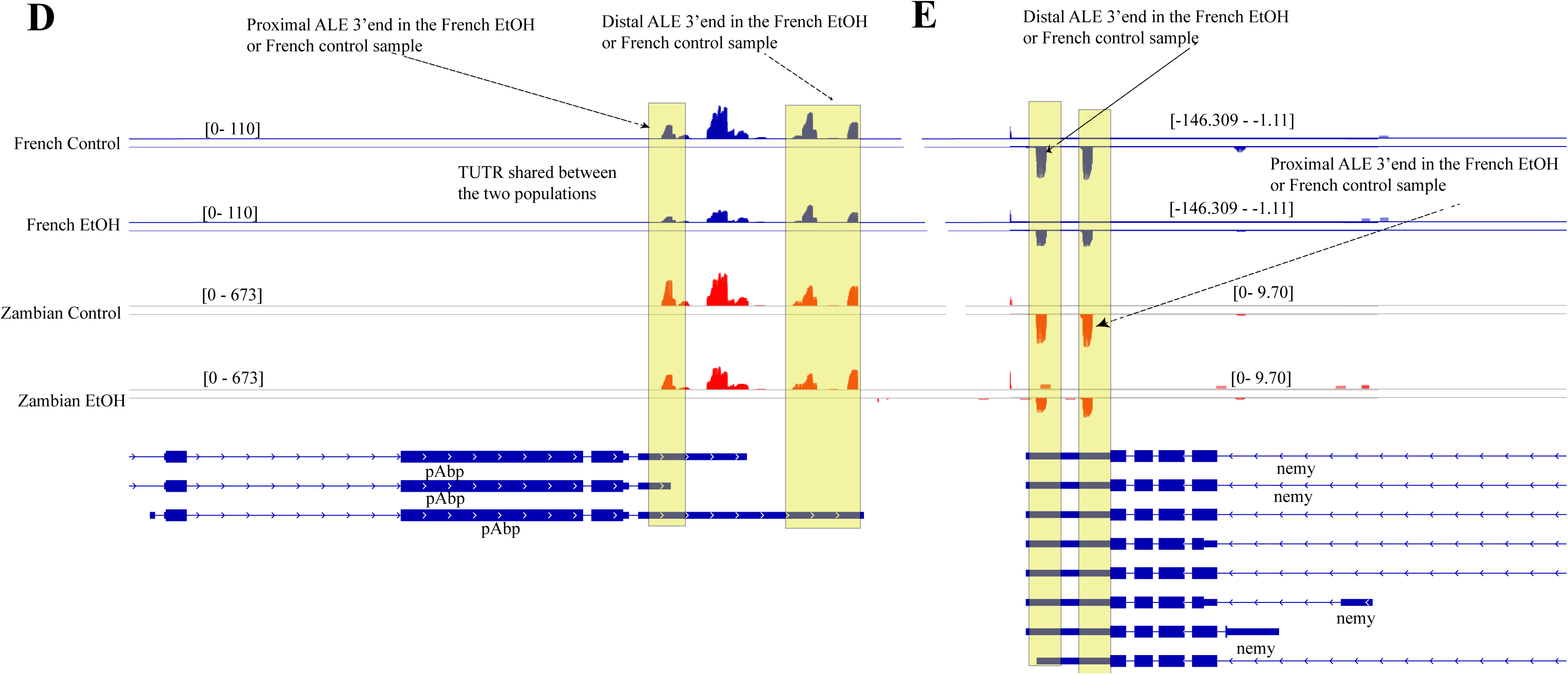
The landscape of APA in both French and Zambian population. **A)** The box plot illustrating the distance between two APA sites across all genotypes. **B)** A bar plot illustrating the number of significant genes per genotype. **C)** Distribution of the top 50 genes ranked by absolute ΔΨ values, illustrating the magnitude and direction of APA regulation in response to ethanol exposure. Each point represents a significant gene (FDR ≤ 0.05, exactly 2 APA sites ≥ 25 nt apart). Genes are plotted on the y-axis, sorted by ΔΨ, while the x-axis represents the ΔΨ value for each gene, defined as Ψ(control)- Ψ(ethanol-treated). Thus, positive ΔΨ values reflect distal site usage in ethanol-treated samples, while negative values reflect proximal site usage. The point shape represent genotype across both French and Zambian population. The size of each dot indicates the APA distance (in nucleotides) between annotated APA sites, offering insight into the scale of transcript lengthening -ranging from modest UTR trimming to substantial isoform changes. **D**) IGV screenshot showing TUTR APA event in *poly(A)-binding protein* and **E**) IGV screenshot of TUTR event in *nemy*. In both examples, reduced read coverage downstream of the proximal poly(A) site in ethanol-treated samples indicates increased proximal site usage. In the IGV screenshots, blues tracks show read coverage for the French sample (control above, ethanol-treated below), while red tracks show read coverage for the Zambian samples in the same. with the blue track underneath, it represents the read coverage for the treated sample. The red tracks represent read coverage for the Zambian population, shown in red. Notable, in both populations, read overage is reduced in the ethanol-treated condition relative to control.

We noticed that by event counts, the French genotypes are enriched for proximal increasing events while the Zambian genotypes are more distal increasing events (Figure 4B). We notice that the magnitude of change/effect size are modest on average, but their dispersion differs by population. For example, in the French population, proximal increasing events are broader in *FR109*/*FR112* while in the Zambian genotypes distal increasing events are broader (Figure 4C).

Consistent with Figure 4B-C, our analysis revealed that across populations, inter-poly(A) distances did not predict whether ethanol drove proximal/distal increasing (France (OR) per distance was approximately = 1.02, *p*=0.87; Zambia vs France interaction *p*=0.78). we noticed that the magnitude of ethanol induced increased with spacing. Each doubling in inter-poly(A) distance was associated with a +51 nt larger absolute change in the 3’ UTR length (*p=*6.7×10^-15^). This association (spacing-magnitude relationship) did not differ by population (interaction *p=*0.17). holding space constant, proximal increasing events were on average +28nt larger than distal (*p*= 0.047). Thus, while spacing does not determine the direction of the ethanol response, it sets the capacity for how long the 3’ UTR change can be. This capacity effect is shared across all population and given the same spacing, proximal increasing events show modestly larger length changes.

### Shared directional APA changes across populations

Genes that exhibit a shared APA shift across the two populations could be considered part of a conserved response to ethanol treatment. We identified 21 genes that were significantly regulated in both the French and Zambian populations and exhibited the same direction of APA change. These genes include APA targets such as *no extended memory, Ubiquitination factor E4B, Ecdysone-induced protein 63F-1, Major Facilitator Superfamily 14 transporter,* and *poly(A)-binding protein*. Their activities span very different molecular functions, although they all sit at regulatory chokepoints that help the cell adapt rapidly to stress or developmental signals. For instance, *no extended memory* tunes neuropeptide maturation in secretory vesicles, affecting neuronal signaling while*, Ubiquitination factor E4B* encodes ubiquitination factor E4B, an E4-type ubiquitin ligase that extends polyubiquitin chains on substrates. *Ube4B* participates in protein polyubiquitination and the ER-associated degradation (ERAD) pathway, contributing to cytoplasmic and nuclear protein quality control and poly(A)-binding protein (*PABP*), is an RNA-binding protein that coats the poly(A) tail of mRNAs to regulate their stability and translation. *pAbp* interacts with translation initiation factors (e.g. *eIF4G*) and has roles in oogenesis, circadian rhythm maintenance in pacemaker neurons, and synaptic function at the neuromuscular junction.

Of the 16 shared tandem UTR events, 10 shifts distally and 6 shift proximally. The 3 shared alternative UTR events include 1 distal shift and 2 proximal shifts. Both shared mixed UTR events lengthen via distal APA. In total, 13 genes underwent distal shifts while 8 underwent proximal shifts. Furthermore, we also noticed that the genomic distances between alternative poly(A) sites differ by direction. Thus, distal events occur on average 492 nt apart (median 291 nt), whereas proximal events span 1077 nt on average (median 803 nt).

### Private changes in response to ethanol

To determine whether 3’ UTR changes in response to ethanol exposure is private to population or genotype, we analyzed the subset of significant APA events that were unique to either the French or Zambian populations. A total of 254 genes exhibited private APA changes, with 159 of them exclusive to the French population and 95 unique to the Zambian population. The distribution of proximal (87) and distal (72) APA events in the French population were somewhat biased towards shortening, while in the Zambian populations there was a more pronounced bias towards distal (64) compared to proximal (31) events. Ethanol based adaptation to APA therefore appears to involve distinct classes of genes between populations and may be characterized by either the suppression of longer isoforms or a bias towards short isoforms. The trend towards tandem UTRs as the primary mode of 3’ UTR regulation held in these private APA events between populations. The French (Zambian) population had more (fewer) diverse types of events, with 123 (79) tandem UTRs, 15 (4) alternative UTRs, and 21 (12) mixed UTR events. This may reflect greater heterogeneity in 3’ UTR usage after ethanol exposure in the French population that is adapted to ethanol.

To assess overall magnitude of change in 3’ UTR length and the extent of potential 3’ UTR APA remodeling in the French and Zambian populations, we compared the shift in 3’ UTR distances between their respective groups. In the French population, distal APA events had a mean poly(A) distance of 978 nt (median of 441 nt), while proximal APA events averaged 1031 nt (median of 374 nt). In contrast, the Zambian population exhibited significantly shorter distances. We noted a distal event average of 832 nt (median of 279 nt), and proximal events averaged 549 nt (median of 264 nt). Thus, the French population recorded a higher number of APA regulated genes and demonstrated a larger magnitude of APA site spacing, indicating more extensive transcript remodeling per gene.

### 3’ UTR changes in opposite directions

Genes that utilize different APA sites after ethanol exposure and show population specific regulation could be core to the adaptive response to ethanol. To determine whether APA regulation differs between the French and Zambian populations, we identified genes that were significantly regulated in both populations but shifted APA usage in different directions in response to ethanol exposure. A total of 21 genes were detected as shared, indicating that APA can proceed through divergent regulatory mechanisms depending on genetic background. These genes include tandem UTR-type events such as *nervana* 1, which favors proximal APA usage in the French population, but shifts toward distal APA usage in Zambian samples (Figure 3D). Other genes showing similar regulatory divergence include *polyadenylation binding protein* (Figure 3E), each demonstrating population specific APA directionality.

### Overlap with existing datasets

Among the French population, we found 53 genes that are jointly regulated by APA and alternative 3’splice sites,^57^ whereas no overlap was detected in the Zambian population. The French overlap genes were enriched for neuronal, and stress responds factors that makes sense biologically in the context of ethanol exposure. The genes include *nervana 3* which is important for sodium pump and ion transport, *vacuolar H^+^-ATPase subunit SFD* which functions as a synaptic vesicle, supporting membrane trafficking and signaling and *no extended memory,* a cytochrome b561-family like to amine systems and memory

## Discussion

Alternative polyadenylation is a widespread post transcriptional mechanism for gene regulation that leads to the generation of mRNA isoforms with different 3’ UTR lengths. In this study, we explored how ethanol affects APA in two genetically distinct *Drosophila melanogaster* populations (France and Zambian) and whether responses depend on population background and genotype. We found that APA were widespread but not uniform. The set of responsive genes, the direction of change (proximal vs distal), the magnitude of 3’ UTR remodeling all varied by population and genotype. These patterns suggests that 3’ UTR regulation via APA is a flexible and strongly genetic-background dependent.

Using a generalized linear model on the event counts (genotypes nested within population), we detected significant population effect and within-population genotype differences. We revealed that the French genotypes show a preference or enrichment for proximal increasing events, while the Zambian genotypes are enriched for distal increasing events. Importantly, these population differences were not explained by APA composition (tandem UTR, alternative UTR, mixed UTR), suggesting that the directional biases reflect broader regulatory context rather than event type differences alone.

Furthermore, for two site genes, the poly(A) spacing spanned a wide range a d acted primarily as a constraint on the remodeling magnitude rather than a determinant of shift direction. Spacing did not predict whether ethanol favored proximal versus distal usage, but larger separation supported larger absolute 3’ UTR length changes, and this scaling was similar across both populations. At comparable scaling, proximal increasing events exhibited modestly larger length changes than distal-increasing events, consistent with spacing setting the capacity of the response while direction is specified by other genotype- and context dependent factors.

Additionally, most APA changes were private to either the French or the Zambian population consistent with genotype-specific regulation. Among private events, French genotypes contained more proximal increasing changes, while the Zambian genotypes contained more distal changes. Tandem UTR events predominated in both cases, with smaller contributions from alternative UTR and mixed UTR. Shared events were relatively few and often differed in direction between populations, suggesting that ethanol responsive APA frequently depends on genetic background. In cases where opposite APA changes were observed (such as in *no extended memory, Major Facilitator Superfamily transporter 14*, *Poly(A)-binding protein, Ecdysone-induced protein 63F-1, Ubiquitination factor E4B)*, the implicated functions (protein quality control and neuronal signaling) suggest plausible phenotypic axes for divergence ^59^.

In addition, considering, *Calmodulin* stood out as the only gene that was significant in all three French genotypes with four APA sites points to a shared ethanol response within the French genotype. Given it central role in Ca²⁺/calmodulin-dependent signaling and prior evidence that CaM/PKA pathways in perineurial glia shape ethanol sensitivity and tolerance APA lengthening ^57^, it may provide a direct post-transcriptional route to tune this signaling axis under ethanol. In this context, the French specific *Calmodulin* signal complements our broader pattern (shortening in *FR112*, balanced in *FR109*, near-zero in *FR113*), suggesting that conserved targets like *Calmodulin* is important for 3’ UTR regulation.

Several mechanisms could contribute to these directional biases. Population differences in the expression or activity of core cleavage and polyadenylation (CPA) factors, RNA-binding proteins that influence tandem UTR selection, or sequence variation near poly(A) signals may all shift the balance between proximal and distal site usage under ethanol exposure. Future work examining cis-regulatory motifs and trans-acting factors will help distinguish among these possibilities.

Together, these results support a model in which genotype-specific APA response with population level directional biases, while the physical distance between polyadenylation sites sets the magnitude of the 3’ UTR remodeling that is achievable. The prevalence of private events and opposite direction changes between the population suggests that natural selection may shape 3’UTR regulation can diverge substantially between genetic backgrounds that differ in ethanol tolerance.

## Acknowledgements

Thanks to John Pool for generously providing the fly strains used in this study, and J. Saltz for feedback on the manuscript.

## Authors’ contributions

SS conceived the study, GBS performed the experiments, bioinformatics analysis, figure generation and drafted the manuscript, SL contributed methods and ECL helped edit the manuscript.

## Availability of data and materials

All data has been made available in the following repositories: The RNA-seq data has been made available under NCBI Bioproject number PRJNA1347849. All data manipulations, statistical tests, and tables were performed in R; figures were created with *ggplot2*(3.5.2)^56^.In addition, all of the relevant code has been deposited at the following link: https://github.com/gsarfo-boateng/3UTR_pipeline Supplementary file 1 contains normalized time series results while Supplementary file 2 contains LABRAT results for the French population while Supplementary file 3 contains results for the Zambian population. Note that the results reflect changes in the ethanol treated sample relative to their controls. Statistical results can be found in Supplementary file 4.

